# MRE11 is crucial for malaria transmission and its absence affects expression of interconnected networks of key genes essential for life

**DOI:** 10.1101/2020.08.24.258657

**Authors:** David S. Guttery, Abhinay Ramaprasad, David J. P. Ferguson, Mohammad Zeeshan, Rajan Pandey, Declan Brady, Anthony A. Holder, Arnab Pain, Rita Tewari

## Abstract

The <u>M</u>eiotic <u>R</u>ecombination 11 protein (MRE11) plays a key role in DNA damage response and maintenance of genome stability. However, little is known about its function during development of the malaria parasite *Plasmodium*. Here, we present a functional, ultrastructural and transcriptomic analysis of *Plasmodium* MRE11 during its life-cycle in both mammalian and mosquito vector hosts. Genetic disruption of *Plasmodium berghei mre11* (PbMRE11) results in significant retardation of oocyst development in the mosquito midgut associated with cytoplasmic and nuclear degeneration, along with concomitant ablation of sporogony and subsequent parasite transmission. Further, absence of PbMRE11 results in significant transcriptional downregulation of genes involved in key interconnected biological processes that are fundamental to all eukaryotic life including ribonucleoprotein biogenesis, spliceosome function and iron-sulphur cluster assembly. Overall, our study provides a comprehensive functional analysis of MRE11’s role in *Plasmodium* development during the mosquito stages and offers a potential target for therapeutic intervention during malaria parasite transmission.

## Introduction

Malaria is one of the deadliest infectious diseases worldwide, for example there were over 219 million clinical infections and 435,000 malaria deaths in 2017 (*1*). Caused by infection with the apicomplexan parasite *Plasmodium* spp., malaria is transmitted by the *Anopheles* mosquito. The parasite lifecycle has several morphologically distinct developmental stages in the mammalian host and mosquito vector (*2*). The host is infected by sporozoites, which migrate in the blood to the liver, where the parasite replicates asexually multiple times in a hepatocyte to produce merozoites. Each merozoite infects a red blood cell, followed by the cycles of intra-erythrocytic multiplication and merozoite release, which are responsible for the clinical symptoms. Some intraerythrocytic parasites become developmentally-arrested gametocytes, which when ingested by the vector initiate the sexual development of the parasite. Gametogenesis and fertilization produce a motile ookinete that invades the mosquito gut wall and forms an oocyst where sporozoites develop and once mature, migrate to the mosquito’s salivary glands for transmission to the host during feeding.

Maintenance of genome stability is fundamental for all organisms and damage to DNA triggers robust DNA damage response (DDR) mechanisms, well-studied in mammalian systems. A signaling pathway is initiated when DNA lesions are identified, which then recruits different DNA repair pathways to repair the damage and preserve genomic integrity. Depending on the level of damage, the DDR pathway can also initiate apoptosis or cell senescence, leading to cell death and limiting further DNA damage due to high genomic instability (*3*). Although the unique molecular aspects of *Plasmodium* DNA replication and cell multiplication are beginning to emerge (*4*), little is known of the key molecular players used to maintain genome stability, especially during DDR. The haploid parasite genome is highly susceptible to extensive DNA damage, such as double-strand breaks (DSBs) resulting from innate DNA replication errors or exposure to radiation and chemical mutagens (*5*). DSBs activate the DSB repair (DSBR) pathway, facilitating repair via two distinct mechanisms: “error-prone” non-homologous end-joining (NHEJ) and “error-free” homologous recombination (HR). In *Plasmodium,* HR appears to be the predominant DDR pathway (*6*), with homologues of many key players of mammalian HR (*5*).

A key initiator of the HR pathway is meiotic recombination 11 protein (MRE11) (*7*), which in mammals is part of the MRN/X complex that includes MRE11, RAD50 and NBS1 (Nijmegen breakage syndrome 1, a homologue of *Saccharomyces cerevisae* Xrs2). This complex has a crucial role in DDR during mitosis, promoting HR between sister chromatids to repair mutations arising during DNA replication (*8*). It acts in two ways: first as an upstream DNA damage sensor (*9*), which activates the ataxia-telangiectasia-mutated (ATM) pathway (*10*), and second to activate DNA repair by promoting bridge formation to allow resection of DSBs (*11*). In parasitic protozoa, MRE11 is not essential in *Trypanosoma brucei* bloodstream forms, but its deletion results in impaired HR, reduced growth and increased sensitivity to DNA double-strand breaks (*12*). Numerous proteins potentially involved in HR have been identified in *P. falciparum* (*5*), although NBS1 is not encoded in the genome. In a recent study, a single *mre11* orthologue containing 2 incomplete but catalytically active MPP_MRE11 domains was identified in *P. falciparum* (Gene ID: PF3D7_0107800) (*15*), and complementation experiments confirmed DDR activity in response to DNA damage (*14*). A role in DDR was confirmed when *P. falciparum mre11* (along with its MRN partner *rad50*) was shown to be significantly upregulated in response to treatment to induce DNA damage with the alkylating agent, methyl methanesulfonate (MMS) (*15*).

The DSBR pathway is a target of intense investigation for the development of cancer therapies (*16*). However, despite the initial studies in *Apicomplexa,* the function of MRE11 throughout the *Plasmodium* lifecycle is still unclear and genome-wide studies have not investigated its role at the different stages (*17, 18*). Here we use the rodent malaria parasite *Plasmodium berghei* (Pb) to provide a functional, ultrastructural and transcriptomic analysis of *Pbmre11*. By gene deletion, we show that MRE11 has an essential role during oocyst maturation and sporogony, which is contributed through the female gamete. We also show by RNA-seq analysis that MRE11’s absence results in significant downregulation of transcription of essential genes that play key roles in ribonucleoprotein biogenesis, spliceosome function and iron-sulfur cluster assembly. These findings suggest that MRE11 has a crucial role during parasite transmission, and may affect vital biological processes through reduced activation of the DNA damage repair response. Overall, our studies offers a potential target for therapeutic intervention of a parasite that still has a huge socioeconomic impact.

## Results

### MRE11-GFP has a nuclear location and is female-cell lineage specific

We first examined the expression and subcellular location of MRE11. *mre11m* RNA is present in cells throughout the *P. berghei* life-cycle (except merozoites), based on single cell RNA-seq data (*19*), with the highest abundance in ookinetes/oocysts (Fig. S1A). To investigate MRE11 protein expression and location throughout the life-cycle, we generated a fusion protein with a C-terminal GFP-tag by single crossover recombination at the endogenous *mre11* locus (PBANKA_020560) (Fig. S1B). Successful integration was confirmed using diagnostic PCR (Fig. S1C) and protein expression confirmed by Western blot using an anti-GFP antibody (Fig. S1D). A ~154 kDa MRE11-GFP was detected in lysates from activated gametocytes, compared with the 29 kDa GFP extracted from a parasite line constitutively expressing GFP (GFPcon 507 cl1 – henceforth called WT-GFP) (*20*).

By live cell imaging of GFP fluorescence, MRE11-GFP was not present in asexual blood stage parasites, but was detected in the first sexual stages, more specifically in the nucleus of female but not male gametocytes (Fig. 1). Following gametocyte activation, female, but not male, gametocytes continued to express the protein. After fertilization the protein remained associated with the nucleus throughout zygote/ookinete development; whereas in oocysts the expression was diffuse and in sporozoites it was focused at a single point adjacent to the nuclear DNA.

**Fig. 1:**
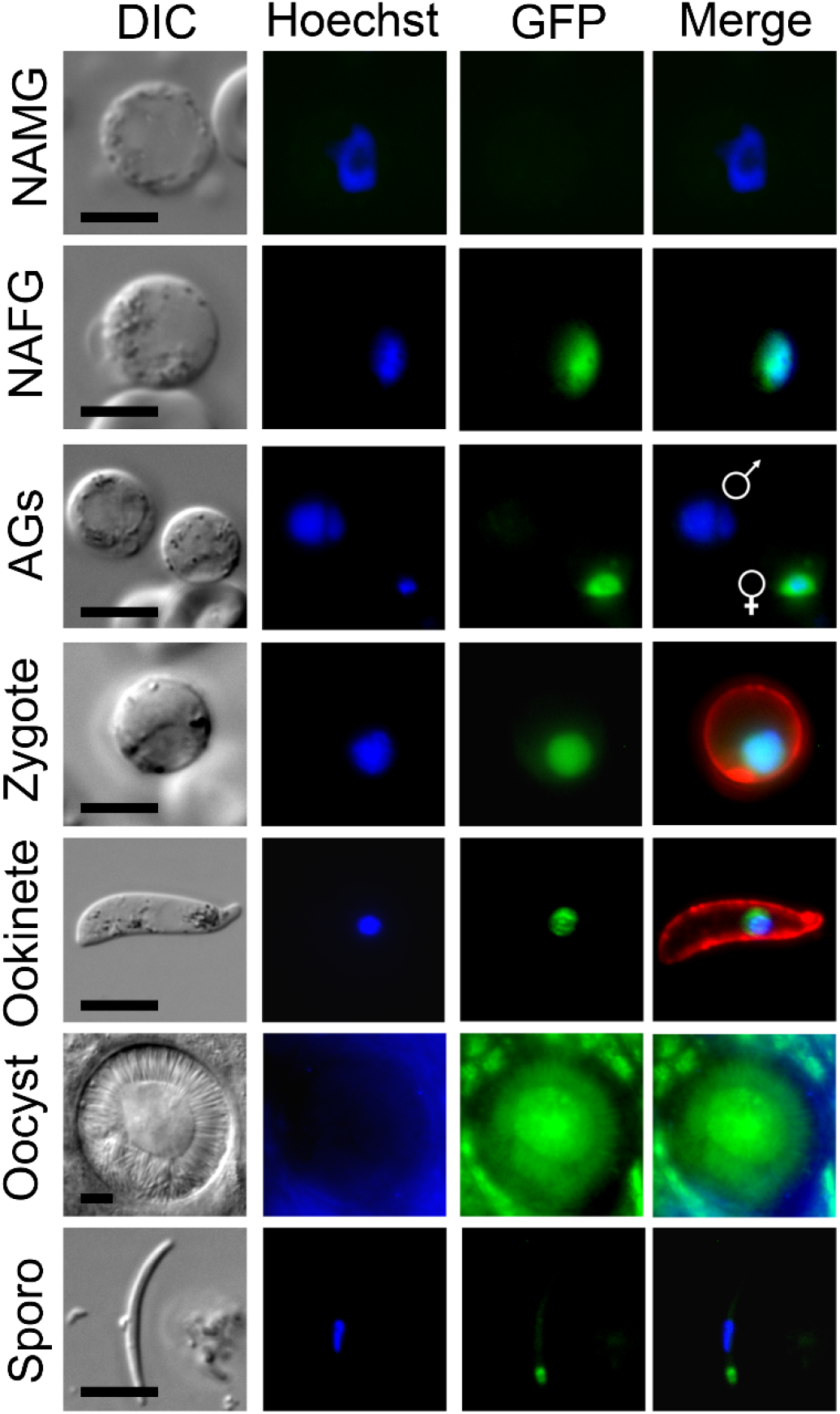
PbMRE11-GFP protein expression throughout most stages of the life cycle. Expression of PbMRE11-GFP in gametocytes, zygotes, ookinetes, oocysts and sporozoites. No GFP expression was observed in asexual ring, trophozoite or schizont stages. NAMG = non-activated male gametocytes; NAFG = non-activated female gametes, AGs = activated gametocytes; Scale bar = 5 μm.

### MRE11 is required for oocyst maturation and sporogony in the mosquito

To determine the function of MRE11 throughout the parasite’s lifecycle, we analyzed asexual blood stages in mice and sexual stages in the mosquito. Using asexual blood stage parasites where the protein is not present, we utilized double crossover homologous recombination at the endogenous *mre11* locus to replace the gene with the *T. gondii dhfr/ts* gene that confers pyrimethamine-resistance. Diagnostic PCR and Southern blotting confirmed successful integration of the targeting construct at the *mre11* locus (Fig. S2). Two independent parasite clones with the *mre11* gene deleted (*mre11* clone 2 and *mre11* clone 9 – henceforward called Δ*mre11*) were analyzed.

WT-GFP control and Δ*mre11* parasites were indistinguishable in asexual blood stage proliferation, microgametocyte exflagellation (Fig. 2A) or ookinete conversion *in vitro* (Fig. 2B), with no significant differences. Similarly, neither ookinete DNA content nor ookinete motility differed in Δ*mre11* lines from WT-GFP controls (Fig. 2C, D and Movie S1, S2). We then investigated whether MRE11 is essential for oocyst development in the mosquito gut. We fed female *A. stephensi* mosquitoes either Δ*mre11* or WT-GFP parasites and analyzed oocyst development and sporogony in the mosquito gut wall at 14 days post-infection. There were significantly fewer oocysts in Δ*mre11* lines (Fig. 2E), and the vast majority of these oocysts were significantly smaller than WT-GFP oocysts (Fig. 2F). In none of the Δ*mre11* oocysts was there any evidence of sporozoite development (Fig. 2G), and those of similar size to WT-GFP oocysts had patterns of fragmented GFP expression and reduced Hoechst DNA-staining. Investigation of the sex-cell lineage of this defect revealed that it could be rescued by crossing Δ*mre11* parasites with Δ*map2* (male-defective) cells (*21*), suggesting that the function of MRE11 is inherited through the female (Fig. 2H).

**Fig. 2:**
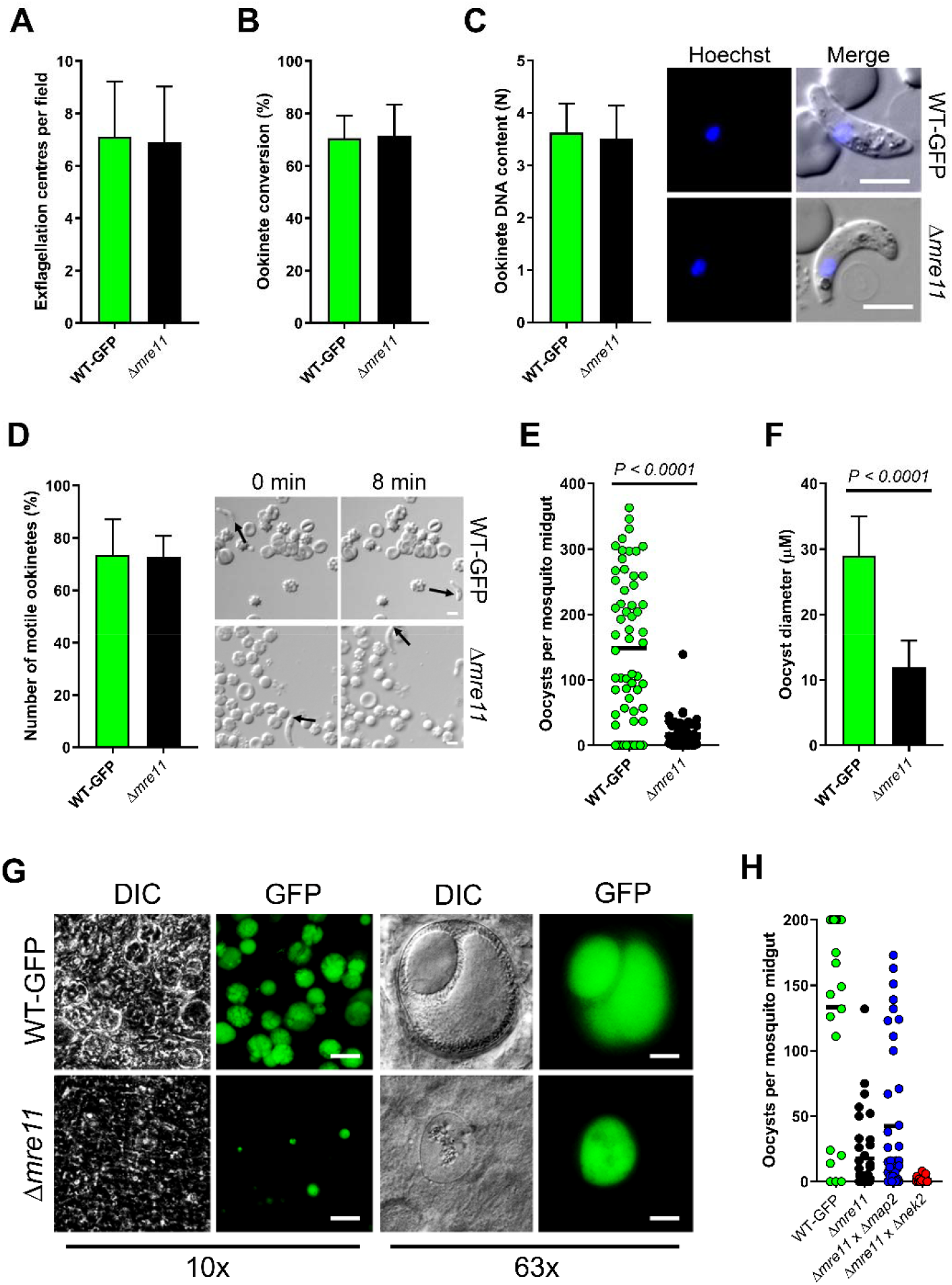
PbMRE11 is dispensable in asexual and sexual stages but important for oocyst development and sporogony. (A) Microgametogenesis of Δ*mre11* lines compared with WT-GFP controls measured as the number of exflagellation centres per field. Means ± SEM are shown. n = 20 from 3 independent experiments. (B) Ookinete conversion as a percentage in Δ*mre11* and WT-GFP lines. Ookinetes were identified using the marker P28 and defined as those cells that successfully differentiated into elongated ‘banana shaped’, stage 6 ookinetes. Bar is the mean ± SEM. n = 3 independent experiments, with >200 cells counted in each. (C) DNA content of mature ookinetes. Bar is the mean ±SD. n = 36 each for WT-GFP and Δ*mre11* lines. (D) Representative frames from time-lapse movies of WT (upper panels) and Δ*mre11* (lower panels) ookinetes in Matrigel. Arrow indicates the apical end of the ookinete. Bar = 10 μm. A number of individual WT or Δ*mre11* ookinetes from 24 hr cultures was measured over 8 min. Bar is the mean ± SEM; n = 3 biological replicates with 3 ookinetes measured in each as technical replicates. (E) Total number of GFP-positive oocysts per infected mosquito, at 14 dpi for Δ*mre11* and WT-GFP lines. Bar = arithmetic mean. n = 3 independent experiments (20 mosquitoes for each). (F) Comparison of oocyst size in Δ*mre11* and WT-GFP lines at 14 dpi. Bar is the mean ±SEM. n = 200 oocysts. (G) Example of relative oocyst size and numbers at 10x and 63x magnification in Δ*mre11* and WT-GFP lines. Images show DIC, Hoechst and GFP at 14 dpi. Scale bar = 50 μm for 10x and 10 μm for 63x. (H) Genetic complementation of Δ*mre11*. Mosquitoes were fed with a combination of WT-GFP, Δ*mre11* or Δ*mre11* with either male (Δ*map2* or female (*Δnek2*) mutants. Shown is a representation of three independent experiments (at least 10 mosquitoes per line, per experiment).

### Ultrastructure analysis reveals degenerative cytoplasm and complete lack of sporozoite formation in Δ*mre11* parasites

To investigate further the defective maturation and sporozoite development of Δ*mre11* parasites, WT-GFP and Δ*mre11* oocysts 14 days post infection were examined by transmission electron microscopy. In WT-GFP lines, numerous large (approximately 70 μm in diameter) oocysts were observed with intact cytoplasm exhibiting various stages of sporozoite formation from early development (Fig. 3A) to some with numerous fully formed sporozoites (Fig. 3B). In contrast, very few Δ*mre11* oocysts were observed and these were much smaller (approximately 20 μm in diameter) than the WT oocysts (compare Fig. 3A, B to Fig. 3C, D). The Δ*mre11* oocysts had a fully formed wall but the cytoplasm exhibited evidence of degeneration (Fig. 3C, D), with vacuolization and organelle breakdown, while the nuclei showed marked swelling of the nuclear membranes (Fig. 3C, D). The degeneration appeared to occur prior to completion of the growth phase and there was no evidence sporozoite formation being initiated.

**Fig. 3:**
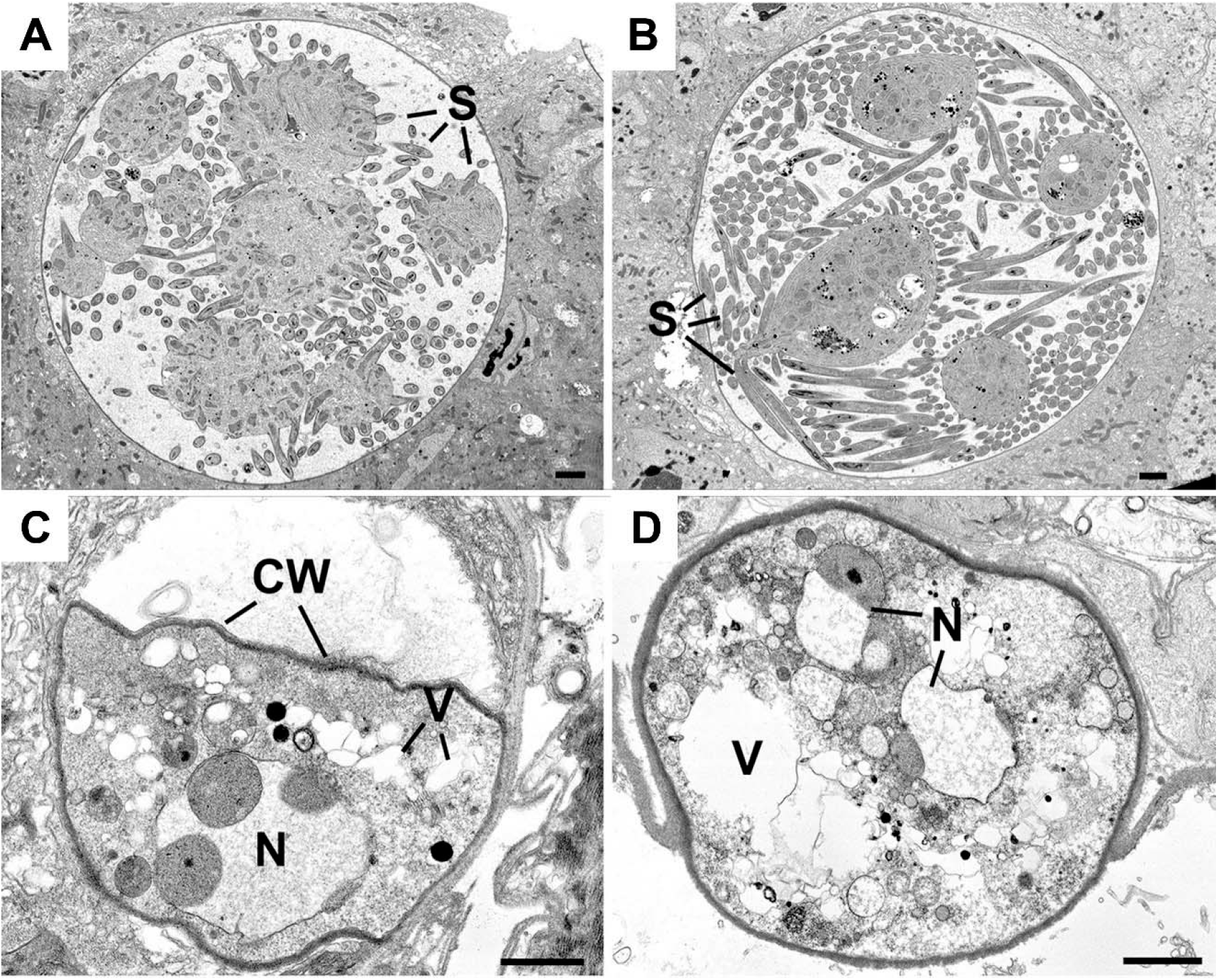
Ultrastructure images of the oocysts of WT and Δ*mre11* parasites at 14 days post-infection. (A) Mid-stage WT oocyst showing early stages of sporozoite formation (S) around the cytoplasmic mass. (B) Late stage WT oocyst showing the large numbers of fully developed sporozoites (S). (C and D) Low power view through the Δ*mre11* oocyst showing the oocyst wall (CW) enclosing the degenerate contents with vacuolated cytoplasm (V) and swollen nucleus (N). Bars represent 10 μm.

### Absence of PbMRE11 results in transcriptional downregulation of key genes essential to all eukaryotic life

To understand better the effect of *mre11-deletion* on the transcriptional footprint of genes across various biological processes, we compared the transcriptomes of Δ*mre11* and WT-GFP parasites by performing RNA-seq on schizonts and a mixture of activated male and female gametocytes. *mre11* transcripts were significantly lowered at both stages in the mutant (by more than 16-fold) compared to those of WT-GFP controls (Fig. 4A), thus confirming loss of *mre11* expression in the knockout parasites. In asexual blood stage schizonts there was no major perturbation of global gene expression resulting from *mre11* deletion, as deduced by comparing the Δ*mre11* and WT-GFP profiles (Fig. 4A and Table S1), which differed only in changes in a small number of genes that mostly belong to highly variable sub-telomeric multigene families. In contrast, there was significant transcriptional dysregulation in activated gametocytes with 418 transcripts (representing 8% of all genes) downregulated in Δ*mre11* parasites (Fig. 4A and Table S1), several of which encode for ribosomal and associated proteins. To define biological signatures within the data, we matched the differentially expressed genes to protein–protein association networks in the STRING database (*22*), and identified major, interconnected gene clusters representing four biological processes: mitochondrial iron-sulfur cluster biogenesis and oxidative DNA damage, ribonucleoprotein (RNP) biogenesis, spliceosome machinery, and host-cell adhesion antigens (Fig. 4B). Of note, downregulated genes involved in mitochondrial iron-sulfur cluster biogenesis and oxidative DNA damage included GLP3, the only glutaredoxin-like protein located in the mitochondrion (*23*), NADP-reductase (*24*), the mitochondrial FAD-linked sulfhydryl oxidase ERV1 (*25*), MMS19-like protein (*26*) and oxoguanine glycosylase 1 (OGG1) (Fig. 4C). Genes essential for RNP biogenesis included those coding for the nuclear chaperone BCP1, the ATPase AKLP1 (an orthologue of hCINAP), nucleolar complex protein 2 (NOC2) and ribonuclease P Protein 1 (RPP1 – depletion of which results in global defects in rRNA processing in *S. cerevisiae* (*27*)). Spliceosome-associated genes included those for debranching enzyme 1 (DBR1) and DBR1 associated ribonuclease 1 (DRN1), microfibril associated protein 1 (MFAP1) and the pre-mRNA-splicing factor CWF7. Finally, genes downregulated in Δ*mre11* gametocytes that are essential for host cell adhesion and parasite transmission in the mosquito included those for secreted ookinete adhesive protein (SOAP), GPI-anchored micronemal antigen (GAMA), gamete release protein (GAMER) and the oocyst markers CAP380 and circumsporozoite protein (CSP) (Fig. 4C). Analysis of single cell RNAseq data (*19*) showed that 85 out of the 99 genes in the four clusters have peak expression during ookinete or oocyst stages (Data file S1), with most of the genes also showing higher levels of transcription in female than male gametocytes (Fig. S1A), further indicating female lineage expression for these genes.

**Fig. 4:**
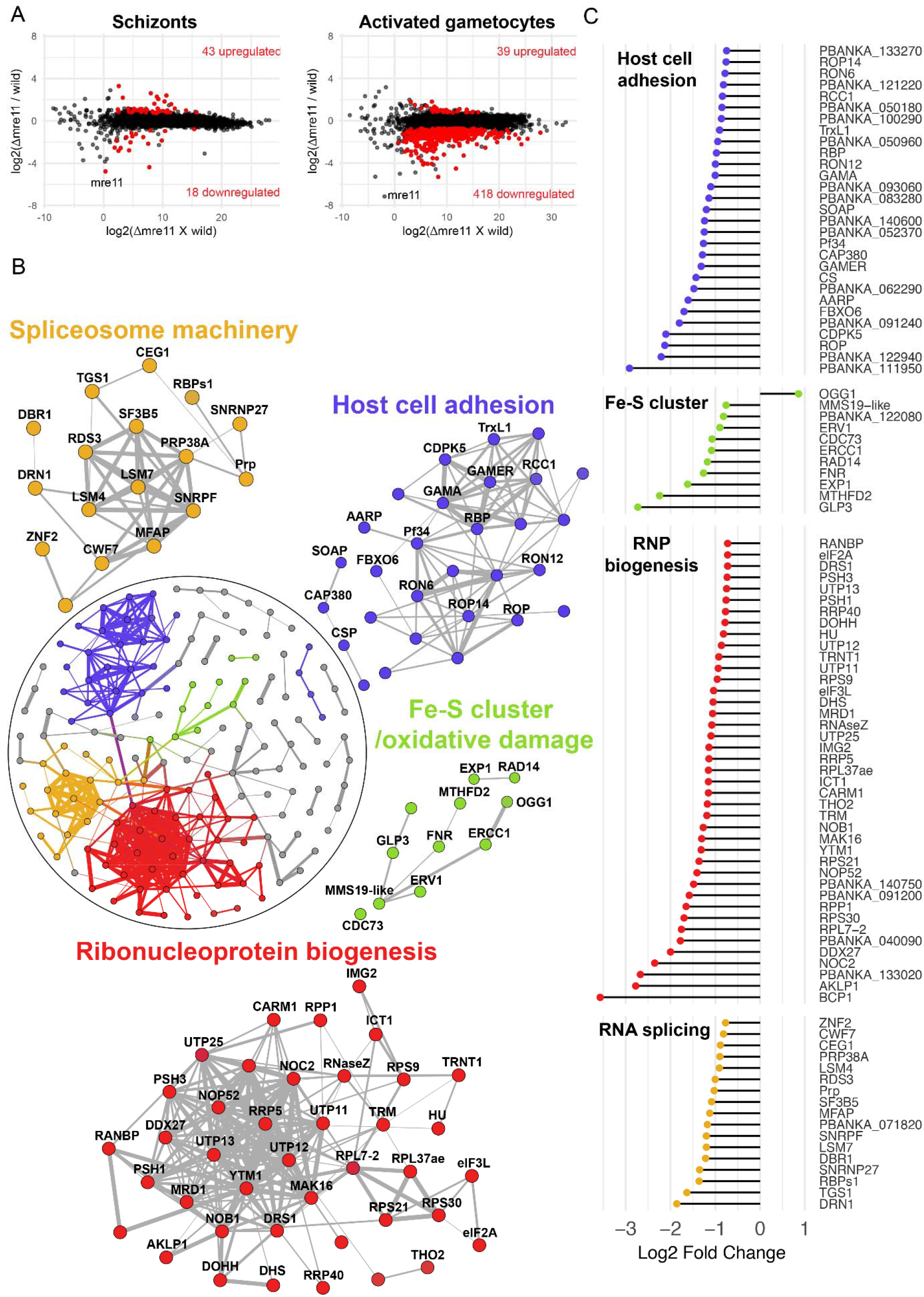
Global transcriptome analysis of Δ*mre11* mutants using RNA-seq. (A) Ratio-intensity plots of quantified FPKM values for each gene in Δ*mre11* and WT-GFP lines at two life stages: schizonts and activated gametocytes. Significant differentially expressed genes (q value<0.05) are in red. Not much perturbation is observed among the Δ*mre11* schizont stage but a significant number of genes are downregulated in Δ*mre11* activated gametocytes. (B) Protein interaction network of differentially expressed genes in Δ*mre11* activated gametocytes. Inset circle shows the complete protein interaction network of 198 proteins (nodes) that had interaction evidence in STRING database (edges). Thickness of the edge denotes confidence of prediction based on several lines of evidence. Four well-interconnected clusters could be determined: ribosome biogenesis (red), spliceosome biogenesis (gold), mitochondria-related iron-sulfur cluster biogenesis and oxidative damage (green) and host cell adhesion molecules (violet). (C) Log2 fold change of genes present in the four main biological processes that were affected in Δ*mre11* lines compared to WT-GFP controls.

## Discussion

Despite its well-known role in mammalian systems for maintenance of genome stability and initiation of DNA damage repair, little is known about MRE11’s functions in *Plasmodium*. Here, we provide a comprehensive functional analysis of its role during parasite sporogony in developing oocysts. Overall, we have shown that MRE11 is a key regulator of malaria parasite transmission and its deletion results in deregulated transcription of essential genes across various key biological processes.

Transcriptional regulation and translational repression of hundreds of genes in female gametocytes are key processes during malaria parasite transmission, occurring in two waves during sporozoite maturation (*28*). Based on MRE11-GFP expression and location, the protein is present only in female gametocytes, and in all subsequent stages of parasite development in the mosquito. We did not detect it in asexual blood stages, which is in contrast to an earlier study showing MRE11 mRNA expression in *P. falciparum* asexual intraerythrocytic stages (*14*), further confirmed by single cell RNA-seq data (*19*). The essential role of translational repression in female gametocytes during parasite transmission to the mosquito (*29, 30*) has been highlighted, but in contrast our MRE11 protein studies suggest that MRE11’s translation is repressed until female gametocyte development. This requirement for the protein in the female lineage is supported by our functional and transcriptomic analyses, which showed that MRE11’s function can be rescued by genetic crossing with a male-defective mutant, and with its enhanced expression in female gametocytes as detected by single-cell RNA-seq (*19*).

Functional analysis showed that MRE11 is not required in asexual blood stages, and its crucial function is during parasite transmission, specifically during oocyst maturation and sporogony. Even though the protein is expressed in the female gametocyte, zygote and ookinete, it is only in the subsequent oocyst stage that a cellular phenotype is observed. Following fertilization, meiosis commences in the ookinete with replication of the diploid DNA. Reductive division, leading finally to haploid sporozoites, presumably occurs in the oocyst, and oocyst development is characterized by 10 or more rounds of DNA replication by closed mitosis and without concomitant cytokinesis to create a syncytial cell (sporoblast) with hundreds of genomes contained within the same nuclear membrane (*31*). MRE11 is known to regulate checkpoint signaling during meiosis (*32*), and since a normal (4N) DNA content was observed in mature ookinetes it is possible that it functions in the reductive division of meiosis in early oocyst development. A previous study analyzing the *Plasmodium* meiotic recombinase <u>D</u>isrupted <u>M</u>eiotic <u>c</u>DNA 1 (DMC1) showed deletion of this genes resulted in a similar phenotype to Δ*mre11* mutants, with significant reduction (up to 80%) in oocyst numbers, which were smaller compared to WT lines and transmission was completely ablated (*33*). However, unlike Δ*mre11* lines sporogony did occur but was slower and limited numbers of nuclei were observed; whereas in Δ*mre11* lines sporogony was completely ablated.

Our global transcriptomic study of Δ*mre11* gametocytes showed that this deletion results in reduced expression of hundreds of genes, many of which are vital for development in all eukaryotes. Ribosome biogenesis is a fundamental component and the primary determinant of translational capacity of the cell (*34*). Formation of functional ribosomes is a complex process that involves transcription, modification and processing of ribosomal RNA, production of ribosomal and auxiliary proteins, and coordinated assembly of ribonucleoprotein (RNP) complexes. Previous ultrastructure analyses of *Plasmodium* oocyst development has shown that the cytoplasm is full of ribosomes and polysomes (*35*); further, female gametocytes are also known to contain higher densities of ribosomes (*36*). Hence, it is likely that the phenotype we observe regarding cytoplasmic degeneration in Δ*mre11* oocysts is caused by reduced transcription of genes essential for RNP biogenesis during these stages. In *Arabidopsis thaliana,* ribosome biogenesis is essential for gametophyte and embryo development and most components are contributed through the female gametophyte lineage (*37*), mirroring our observation of female inheritance of the Δ*mre11* phenotype. An additional key biological network affected in oocysts of Δ*mre11* lines involves spliceosome machinery. Nuclear pre-mRNA splicing is catalyzed by the spliceosome (*38*) and recent findings show that it is a major contributor to genome stability in mammalian cells (*39, 40*). Given that MRE11 is a key modulator of the DNA damage response, our transcriptome data provides further evidence of possible cross-talk (and potential feedback loop) between DNA damage responses, RNA processing/RNP biogenesis and spliceosomes (*41–43*) and show that these complex interrelations are conserved in *Plasmodium*. We also observed significant downregulation of genes associated with iron-sulfur cluster biogenesis and mitochondrial oxidative DNA damage pointing to mitochondrial dysfunction that can contribute to genome instability (*44*). The mitochondrion is the source of reactive oxidative species (ROS) that can cause endogenous DNA damage (*45*) and the possibility of oxidative damage to DNA is shown by an upregulation of OGG1 in Δ*mre11* parasites. The predominant oxidative damage due to ROS is to guanine yielding 7,8-dihydro8-oxoguanine (8-oxoG) and is primarily repaired by OGG1 through the base-excision repair (BER) pathway (*46*). OGG1 is bifunctional and can both remove the damaged base to form an abasic site and also cause a 3’-blocked single-strand break that is then repaired by either AP endonucleases or RAD1/RAD10 mediated nucleotide excision repair (NER) (*47*). It has also been shown previously that OGG1-mediated single-strand breaks can exacerbate oxidative stress-mediated cell death when DNA repair pathways are compromised (*48, 49*). Thus the increase in OGG1 expression coupled with reduction in RAD10, RAD14 and MMS19-like protein gene expression (all playing key roles in the NER pathway) could all contribute to accumulation of deleterious DNA damage that is usually regulated by MRE11. Finally, several studies have highlighted a number of genes that are essential for *Plasmodium* ookinete adhesion and invasion, and oocyst development (*50*), a number of which may be affected due to MRE11’s absence. A previous genome-wide screening study did not analyze MRE11, but it did suggest that RAD50, the MRN complex partner of MRE11, is essential for parasite transmission (*18*). This suggests that MRE11 and RAD50 may form a complex that is essential for successful oocyst development. *MRE11* is important for spore wall formation and in mediating transcriptional regulation during sporulation in *S. cerevisiae* (*51*), consistent with the idea that the crucial function of MRE11 may affect transcriptional regulation during oocyst development. Complementing this, our ultrastructure analysis was consistent with the reduced number of oocysts and, while the oocyst wall formed, there was limited growth and rapid degeneration of the cytoplasm and nucleus. The changes are similar to those exhibited by other mutants that result in oocyst degeneration, although this occurs at an earlier stage before parasite growth (*52*).

While we did not analyze the genome for alterations due to MRE11’s absence, we have shown that a specific downregulation of several vital biological processes interrelated with maintaining genome stability occurs as a result. Taken together, our study provides molecular evidence of the crucial role for MRE11 in regulating sexual stage development and mosquito transmission. Due to its essentiality during mosquito infection, we provide a new potential target for therapeutic intervention against a disease that still has a devastating socioeconomic impact. Further studies on the function of MRE11, such as a role in DNA damage response, will highlight its potential as a therapeutic target.

## Materials and Methods

### Ethics

All animal work has passed an ethical review process in accordance with the United Kingdom ‘Animals (Scientific Procedures) Act 1986’ and in compliance with ‘European Directive 86/609/EEC’ for the protection of animals used for experimental purposes. The project licence number is 40/3344.

### Animals

Six to eight-week old female Tuck-Ordinary (TO) outbred mice (Harlan) were used for all experiments.

### Generation of transgenic parasites

For C-terminal GFP-tagging of MRE11 by single homologous recombination, a 1657 bp region of *mre11* (PBANKA_020560) starting 1494 bp downstream of the ATG start codon and omitting the stop codon was amplified using primers T0751 (5’-CCCCGGTACCCGAAATGAAAGAAAT AGAAGGATTCC-3 ‘) and T0752 (5’-CCCCGGGCCCTTTATTCATTTCTGAAATTGTCGAATTTATA-3’). The DNA fragment was inserted using *KpnI* and *ApaI* restriction sites upstream of the *gfp* sequence in the pOB277 plasmid containing a human *dhfr* cassette to confer resistance to pyrimethamine. The vector was linearized with *BglII* before transfection. The Δ*mre11* gene-knockout targeting vector was constructed using the pBS-DHFR plasmid, which contains polylinker sites flanking a *Toxoplasma gondii dhfr/ts* expression cassette, as described previously (*53*). A 611 bp fragment at the 5’ end of the *mre11* sequence was generated from genomic DNA using PCR primers P0141 (5’-<u>CCCCGGGCCC</u>TTGTGCATACACATCAACAGATAA-3’) and P0142 (5’-<u>GGGGAAGCTT</u>ATCCAAATCTGATAAGTAATTATCCA-3’) and inserted into pBS-DHFR using *Apa*I and *HindIII* restriction sites upstream of the *dhfr/ts* cassette. A 575 bp fragment generated with primers P0143 (5’-<u>CCCCGAATTC</u>GAATGAATTGAAGGATATCCCAG-3’) and P0144 (5’-<u>GGGGTCTAGA</u>CTGTATTGGAGATGAATATTATGGA-3’) from the 3’ region of *mre11* was then inserted downstream of the *dhfr/ts* cassette using *Eco*RI and *XbaI* restriction sites. The linear targeting sequence was released using *ApaI/XbaI* digestion of the plasmid.

*P. berghei* ANKA line 2.34 (for GFP-tagging) or ANKA line 507cl1 (for gene deletion) were transfected by electroporation (*20*). Briefly, electroporated parasites were mixed immediately with 100 μl of reticulocyte-rich blood from a naïve mouse treated with phenylhydrazine (6 mg/ml) (Sigma-Aldrich), incubated at 37°C for 20 min and then injected intraperitoneally into another mouse. From day 1 post infection, 70 μg/ml pyrimethamine (Sigma-Aldrich) was supplied in the drinking water for 4 days. Mice were monitored for the appearance of parasites for fifteen days, then drug-resistant parasites were passaged into a second mouse with continuing drug selection. Parasites were cloned by limiting dilution and genotyped.

### Parasite genotyping and Western blotting

Diagnostic PCR was used with primer 1 (IntP14tag, 5’-CAGATTCACAGATGCATACATA-3’) and primer 2 (ol492) (*54*) to confirm integration of the *mre11* GFP targeting construct. For the gene deletion, diagnostic PCR with primer 3 (IntP14KO, 5’-CCTACGCACCAACTACTCGTTA-3’) and primer 4 (ol248) (*53*) was used to confirm integration of the targeting construct; whereas primer 5 (P014KO1, 5’-AAGATGCAGGAAAAATGAGA-3’) and primer 6 (P014KO2, 5’-GGTTTTGAATTATGCTCGTG-3’) were used to confirm deletion of the *mre11* gene. For Southern blotting, genomic DNA from WT-GFP and mutant parasites was digested with *Hind*III and fractionated on a 0.8% agarose gel before blotting onto a nylon membrane (GE Healthcare). A probe was generated from a PCR fragment homologous to the 3’ sequence just outside of the targeted region using the AlkPhos direct labelling kit (GE Healthcare) according to manufacturer’s instructions. Expected sizes post digestion with *Hind*III were 3 kb and 5 kb for WT-GFP and Δ*mre11* lines, respectively. For confirmation of MRE11-GFP expression, western blot was performed as previously described (*54*).

### RNA-seq transcriptome sequencing

For RNA extraction, parasite samples were passed through a plasmodipur column to remove host DNA contamination prior to RNA isolation. Total RNA was isolated from purified parasites using an RNeasy purification kit (Qiagen). RNA was vacuum concentrated (SpeedVac) and transported using RNA stable tubes (Biomatrica). Total RNA was extracted from schizonts and activated gametocytes of WT-GFP and mutant parasites (three biological replicates each). Strand-specific mRNA sequencing was performed of total RNA and using TruSeq stranded mRNA sample prep kit LT (Illumina), as previously described (*52*). Libraries were sequenced using an Illumina Hiseq with paired-end 100 bp read chemistry. Strand-specific RNA-Seq paired-end reads were mapped onto the *P. berghei* ANKA genome (PlasmoDB v9.2) using TopHat version 2.0.8, quantified and compared across different samples using Cuffdiff version 2.1.1 and statistical analysis performed using cummeRbund package in R. The STRING database version 11 (*22*) was searched for known and predicted protein-protein interactions among the differentially expressed genes. The interaction network was visualized in Gephi (https://gephi.org/) using Fruchterman Reingold layout. Life stage-specific gene expression of certain genes was further explored in the single cell RNA-seq datasets for *P. berghei* published as part of the Malaria Cell Atlas (*19*).

### Phenotypic analysis

Phenotypic analyses were performed as described previously (*55*). Asexual proliferation and gametocytogenesis were analysed by microscopy of blood smears. Gametocyte activation, zygote formation and ookinete conversion rates were assessed by microscopy using in vitro cultures and antibody staining for the surface antigen P28. For ookinete DNA content assays, nuclear fluorescence intensity of WT-GFP or mutant parasites from 24 h cultures stained with Hoechst dye was measured using ImageJ software. Values were expressed relative to the average fluorescence intensity of haploid ring-stage parasites on the same slide and corrected for background fluorescence. Ookinete motility assays used Matrigel (Corning): in brief, ookinete cultures were added to an equal volume of Matrigel on ice, mixed thoroughly, dropped onto a slide, covered with a cover slip, and sealed with nail polish. The Matrigel was then allowed to set at 20°C for 30 min. After identifying a field containing ookinetes, time-lapse videos were recorded at every 5 s for 100 cycles on a Zeiss AxioImager M2 microscope fitted with an AxioCam ICc1 digital camera (Carl Zeiss, Inc). For mosquito transmission, triplicate sets of 20– 60 *Anopheles stephensi* were used. Genetic cross experiments were performed as previously described (*52*).

### Electron microscopy

Mosquito midguts at 14 days post infection (dpi) were fixed in 4% glutaraldehyde in 0.1 M phosphate buffer and processed for electron microscopy as previously described (*54*). Briefly, samples were post-fixed in osmium tetroxide, treated *en bloc* with uranyl acetate, dehydrated and then embedded in Spurr’s epoxy resin. Thin sections were stained with uranyl acetate and lead citrate prior to examination in a JEOL1200EX electron microscope (Jeol UK Ltd).

### Statistical analyses

Statistical analyses were performed using GraphPad Prism 7 (GraphPad Software). For ookinete development, two-way ANOVA was used. For oocyst development, Student’s t-test was used.

## General

We thank Julie Rogers for technical assistance and the personnel at the Bioscience Core Laboratory (BCL) in KAUST for sequencing the RNA samples and producing the raw datasets.

## Funding

This project was funded by MRC Investigator Award and MRC project grants to R.T. [G0900109, G0900278, MR/K011782/1] and also supported by a faculty baseline funding [BAS/1/1020-01-01] to AP. A.A.H. is supported by the Francis Crick Institute (FC010097) which receives its core funding from Cancer Research UK (FC010097), the UK Medical Research Council (FC010097). A.P. and A.R. are funded by KAUST. The funders had no role in study design, data collection and analysis, decision to publish, or preparation of the manuscript.

## Author contributions

Conceived and designed the experiments: RT, DSG and AAH. Performed the experiments: RT, AR, RP, DJPF, AP, MZ, DB and DSG. Analyzed the data: DSG, AR, RP, RT, AAH, MZ, DB, DJPF, AP and DSG. Contributed reagents/materials/analysis tools: DJPF, AAH, AR, AP and RT. Wrote the paper: DSG, AR, DJPF, AP, AAH and RT. Performed the functional and GFP tagging experiments: DSG, MZ, DB, RP and RT. AR and DG performed database searches, sequence-based analysis, and other bioinformatics analysis. Electron microscopy experiments: DJPF. RNA-seq and data analysis was performed by AR and AP.

## Competing interests

The authors declare that no competing interests exist.

## Data and materials availability

The raw RNA sequencing data has been deposited in the European Nucleotide Archive (https://www.ebi.ac.uk/ena) through accession numbers-ERR409988, ERR409989, ERR409990, ERR409994, ERR409995 and ERR409996.

## Notes

### Competing Interest Statement

The authors have declared no competing interest.

## References and Notes

1. W. H. Organisation, “World Malaria Report 2018,” (World Health Organization, 2018).

2. L. H. Bannister, I. W. Sherman, in eLS. (John Wiley & Sons, Ltd, 2001).

3. A. Ciccia, S. J. Elledge, The DNA damage response: making it safe to play with knives. Mol Cell 40, 179–204 (2010).

4. D. S. Guttery, M. Roques, A. A. Holder, R. Tewari, Commit and Transmit: Molecular Players in Plasmodium Sexual Development and Zygote Differentiation. Trends Parasitol 31, 676–685 (2015).

5. A. H. Lee, L. S. Symington, D. A. Fidock, DNA repair mechanisms and their biological roles in the malaria parasite Plasmodium falciparum. Microbiol Mol Biol Rev 78, 469–486 (2014).

6. L. A. Kirkman, E. A. Lawrence, K. W. Deitsch, Malaria parasites utilize both homologous recombination and alternative end joining pathways to maintain genome integrity. Nucleic acids research 42, 370–379 (2014).

7. T. H. Stracker, J. H. Petrini, The MRE11 complex: starting from the ends. Nat Rev Mol Cell Biol 12, 90–103 (2011).

8. D. A. Bressan, B. K. Baxter, J. H. Petrini, The Mre11-Rad50-Xrs2 protein complex facilitates homologous recombination-based double-strand break repair in Saccharomyces cerevisiae. Mol Cell Biol 19, 7681–7687 (1999).

9. M. F. Lavin, ATM and the Mre11 complex combine to recognize and signal DNA double-strand breaks. Oncogene 26, 7749–7758 (2007).

10. Y. Shiloh, Y. Ziv, The ATM protein kinase: regulating the cellular response to genotoxic stress, and more. Nat Rev Mol Cell Biol 14, 197–210 (2013).

11. L. S. Symington, Mechanism and regulation of DNA end resection in eukaryotes. Crit Rev Biochem Mol Biol 51, 195–212 (2016).

12. K. S. Tan, S. T. Leal, G. A. Cross, Trypanosoma brucei MRE11 is non-essential but influences growth, homologous recombination and DNA double-strand break repair. Molecular and biochemical parasitology 125, 11–21 (2002).

13. R. Pandey et al., Genome wide in silico analysis of Plasmodium falciparum phosphatome. BMC Genomics 15, 1024 (2014).

14. S. B. Badugu et al., Identification of Plasmodium falciparum DNA Repair Protein Mre11 with an Evolutionarily Conserved Nuclease Function. PloS one 10, e0125358 (2015).

15. D. K. Gupta, A. T. Patra, L. Zhu, A. P. Gupta, Z. Bozdech, DNA damage regulation and its role in drug-related phenotypes in the malaria parasites. Sci Rep 6, 23603 (2016).

16. M. J. O’Connor, Targeting the DNA Damage Response in Cancer. Mol Cell 60, 547–560 (2015).

17. E. Bushell et al., Functional Profiling of a Plasmodium Genome Reveals an Abundance of Essential Genes. Cell 170, 260–272 e268 (2017).

18. R. R. Stanway et al., Genome-Scale Identification of Essential Metabolic Processes for Targeting the Plasmodium Liver Stage. Cell 179, 1112–1128 e1126 (2019).

19. V. M. Howick et al., The Malaria Cell Atlas: Single parasite transcriptomes across the complete Plasmodium life cycle. Science 365, (2019).

20. C. J. Janse et al., High efficiency transfection of Plasmodium berghei facilitates novel selection procedures. Molecular and biochemical parasitology 145, 60–70 (2006).

21. R. Tewari, D. Dorin, R. Moon, C. Doerig, O. Billker, An atypical mitogen-activated protein kinase controls cytokinesis and flagellar motility during male gamete formation in a malaria parasite. Molecular microbiology 58, 1253–1263 (2005).

22. D. Szklarczyk et al., STRING v11: protein-protein association networks with increased coverage, supporting functional discovery in genome-wide experimental datasets. Nucleic acids research 47, D607–D613 (2019).

23. S. Kehr, N. Sturm, S. Rahlfs, J. M. Przyborski, K. Becker, Compartmentation of redox metabolism in malaria parasites. PLoS Pathog 6, e1001242 (2010).

24. R. Yan, S. Adinolfi, A. Pastore, Ferredoxin, in conjunction with NADPH and ferredoxin-NADP reductase, transfers electrons to the IscS/IscU complex to promote iron-sulfur cluster assembly. Biochim Biophys Acta 1854, 1113–1117 (2015).

25. M. Rissler et al., The essential mitochondrial protein Erv1 cooperates with Mia40 in biogenesis of intermembrane space proteins. J Mol Biol 353, 485–492 (2005).

26. O. Stehling et al., MMS19 assembles iron-sulfur proteins required for DNA metabolism and genomic integrity. Science 337, 195–199 (2012).

27. V. Stolc, S. Altman, Rpp1, an essential protein subunit of nuclear RNase P required for processing of precursor tRNA and 35S precursor rRNA in Saccharomyces cerevisiae. Genes Dev 11, 2926–2937 (1997).

28. S. E. Lindner et al., Transcriptomics and proteomics reveal two waves of translational repression during the maturation of malaria parasite sporozoites. Nat Commun 10, 4964 (2019).

29. G. R. Mair et al., Regulation of sexual development of Plasmodium by translational repression. Science 313, 667–669 (2006).

30. G. R. Mair et al., Universal features of post-transcriptional gene regulation are critical for Plasmodium zygote development. PLoS Pathog 6, e1000767 (2010).

31. H. Matthews, C. W. Duffy, C. J. Merrick, Checks and balances? DNA replication and the cell cycle in Plasmodium. Parasit Vectors 11, 216 (2018).

32. V. Borde, The multiple roles of the Mre11 complex for meiotic recombination. Chromosome Res 15, 551–563 (2007).

33. G. Mlambo, I. Coppens, N. Kumar, Aberrant sporogonic development of Dmc1 (a meiotic recombinase) deficient Plasmodium berghei parasites. PloS one 7, e52480 (2012).

34. E. Thomson, S. Ferreira-Cerca, E. Hurt, Eukaryotic ribosome biogenesis at a glance. J Cell Sci 126, 4815–4821 (2013).

35. E. U. Canning, R. E. Sinden, The organization of the ookinete and observations on nuclear division in oocysts of Plasmodium berghei. Parasitology 67, 29–40 (1973).

36. D. A. Baker, Malaria gametocytogenesis. Molecular and biochemical parasitology 172, 57–65 (2010).

37. S. Missbach et al., 40S ribosome biogenesis co-factors are essential for gametophyte and embryo development. PloS one 8, e54084 (2013).

38. C. L. Will, R. Luhrmann, Spliceosome structure and function. Cold Spring Harb Perspect Biol 3, (2011).

39. B. Adamson, A. Smogorzewska, F. D. Sigoillot, R. W. King, S. J. Elledge, A genome-wide homologous recombination screen identifies the RNA-binding protein RBMX as a component of the DNA-damage response. Nat Cell Biol 14, 318–328 (2012).

40. M. Tanikawa, K. Sanjiv, T. Helleday, P. Herr, O. Mortusewicz, The spliceosome U2 snRNP factors promote genome stability through distinct mechanisms; transcription of repair factors and R-loop processing. Oncogenesis 5, e280 (2016).

41. C. Naro, P. Bielli, V. Pagliarini, C. Sette, The interplay between DNA damage response and RNA processing: the unexpected role of splicing factors as gatekeepers of genome stability. Front Genet 6, 142 (2015).

42. A. S. Bader, B. R. Hawley, A. Wilczynska, M. Bushell, The roles of RNA in DNA double-strand break repair. Br J Cancer 122, 613–623 (2020).

43. V. O. Wickramasinghe, A. R. Venkitaraman, RNA Processing and Genome Stability: Cause and Consequence. Mol Cell 61, 496–505 (2016).

44. J. R. Veatch, M. A. McMurray, Z. W. Nelson, D. E. Gottschling, Mitochondrial dysfunction leads to nuclear genome instability via an iron-sulfur cluster defect. Cell 137, 1247–1258 (2009).

45. L. A. Sena, N. S. Chandel, Physiological roles of mitochondrial reactive oxygen species. Mol Cell 48, 158–167 (2012).

46. X. Zhou et al., OGG1 is essential in oxidative stress induced DNA demethylation. Cell Signal 28, 1163–1171 (2016).

47. A. D. Scott et al., Spontaneous mutation, oxidative DNA damage, and the roles of base and nucleotide excision repair in the yeast Saccharomyces cerevisiae. Yeast 15, 205–218 (1999).

48. S. N. Guzder et al., Requirement of yeast Rad1-Rad10 nuclease for the removal of 3’-blocked termini from DNA strand breaks induced by reactive oxygen species. Genes Dev 18, 2283–2291 (2004).

49. M. Guillet, S. Boiteux, Endogenous DNA abasic sites cause cell death in the absence of Apn1, Apn2 and Rad1/Rad10 in Saccharomyces cerevisiae. EMBO J 21, 2833–2841 (2002).

50. R. C. Smith, C. Barillas-Mury, Plasmodium Oocysts: Overlooked Targets of Mosquito Immunity. Trends Parasitol 32, 979–990 (2016).

51. K. Kugou, H. Sasanuma, K. Matsumoto, K. Shirahige, K. Ohta, Mre11 mediates gene regulation in yeast spore development. Genes Genet Syst 82, 21–33 (2007).

52. M. Roques et al., Plasmodium P-Type Cyclin CYC3 Modulates Endomitotic Growth during Oocyst Development in Mosquitoes. PLoS Pathog 11, e1005273 (2015).

53. R. Tewari et al., The Systematic Functional Analysis of Plasmodium Protein Kinases Identifies Essential Regulators of Mosquito Transmission. Cell Host & Microbe 8, 377–387 (2010).

54. David S. Guttery et al., Genome-wide Functional Analysis of Plasmodium Protein Phosphatases Reveals Key Regulators of Parasite Development and Differentiation. Cell Host & Microbe 16, 128–140 (2014).

55. E. M. Patzewitz et al., An ancient protein phosphatase, SHLP1, is critical to microneme development in Plasmodium ookinetes and parasite transmission. Cell Rep 3, 622–629 (2013).

