## Supplemental Information for "MRE11 is crucial for malaria transmission and its absence affects expression of interconnected networks of key genes essential for life"

Supplementary Materials

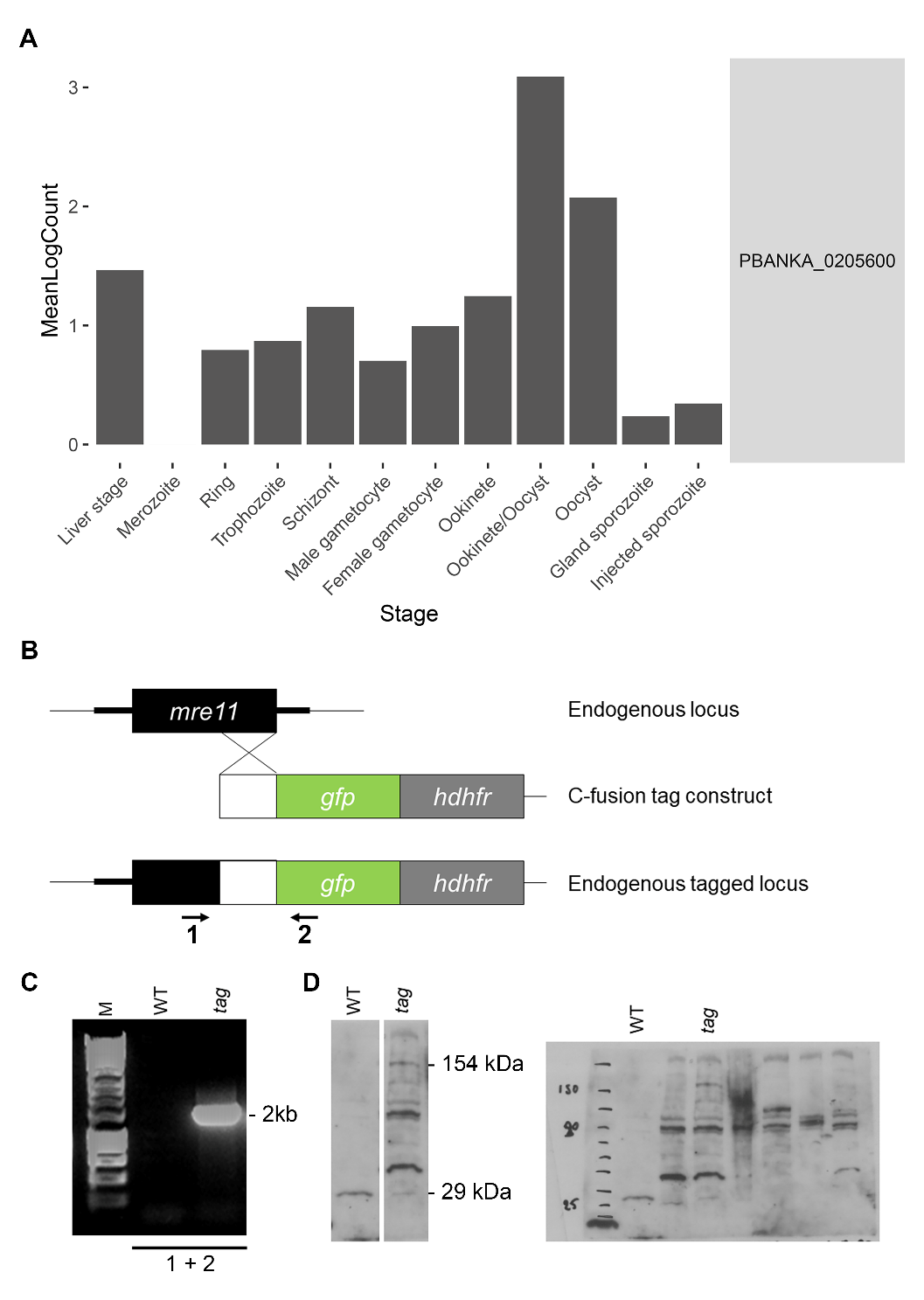

**Fig. S1: GFP-tagging of the endogenous *mre11* locus and *mre11* RNA expression.** (A) Life stage-wise gene expression of *mre11* based on *Plasmodium berghei* single cell RNA-seq data (*18*). (B) Schematic representation of the gene targeting strategy used for gene tagging the endogenous locus with *gfp* via single homologous recombination. Primers 1+2 used for diagnostic PCR are indicated. (C) Diagnostic PCR confirming successful integration of the tagging sequence. (D) Left: Western blot analysis using an anti-GFP antibody against control wild-type-GFP (WT) and transgenic (tag) activated gametocytes showing bands of expected sizes of 29 kDa for wild-type-GFP and ~154 kDa for MRE11-GFP. Right: original Western blot image

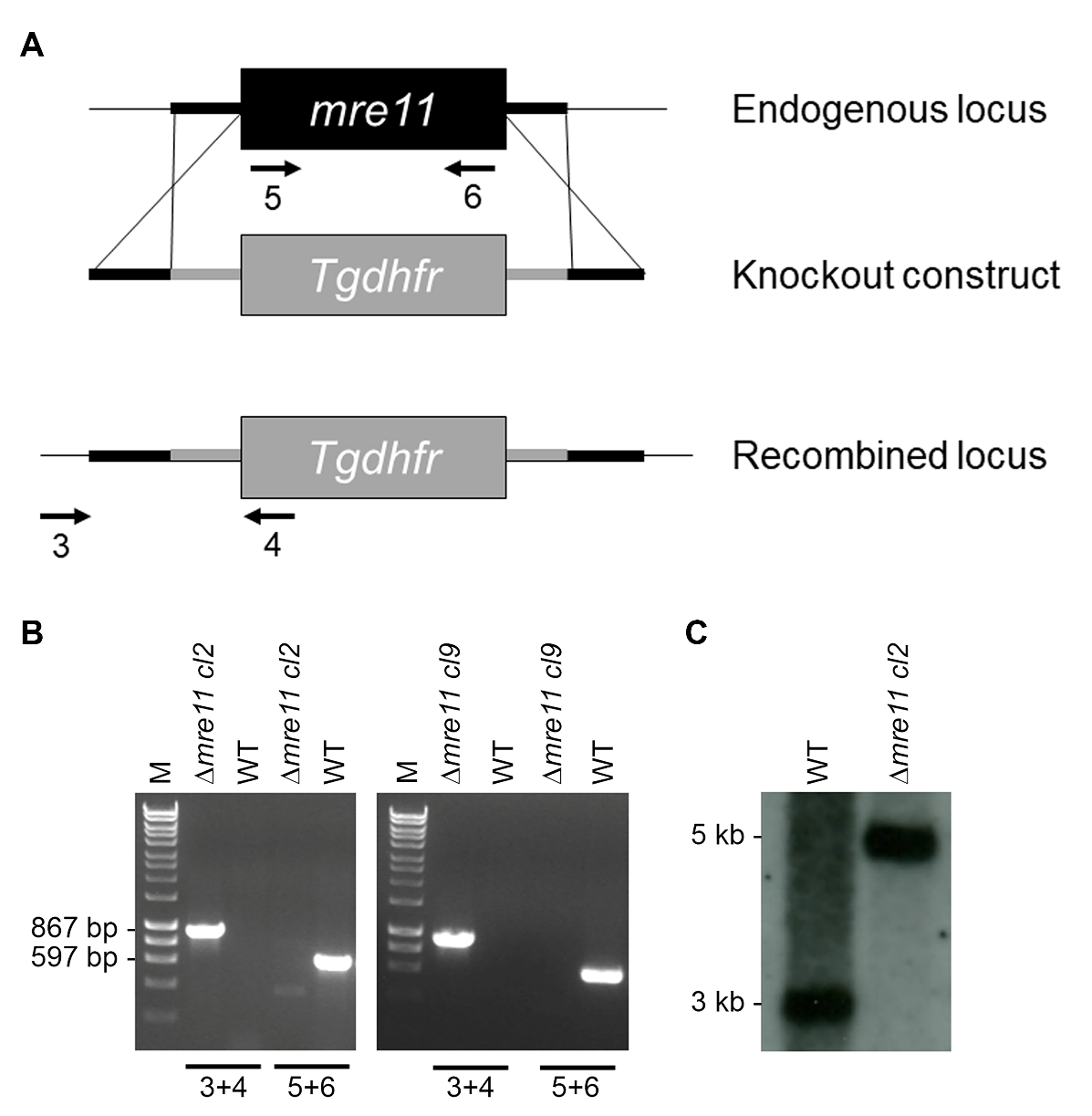

**Fig. S2: Deletion of the *mre11* gene.** (A) Schematic representation of the gene targeting strategy used for gene disruption via double homologous recombination. Primers 3–6 used for diagnostic PCR are indicated. (B) Diagnostic PCR confirming successful integration of the disruption sequence of *mre11* in the *P. berghei* ANKA 2.34 line constitutively expressing GFP. Primers 3+4 were used to verify successful integration at the correct locus. Primers 5+6 were used to confirm loss of the endogenous gene. (C) Southern blot analysis of *Hind*III digested clone 2 genomic DNA using the 3′ UTR of the targeting construct as a probe. Band sizes for Δ*mre11* and wild-type (WT-GFP) are indicated.

**Table S1: RNA-Seq analysis of WT-GFP and Δ*mre11* lines.** Data shows differential expression of global transcripts significantly deregulated in Δ*mre11* lines compared to WT-GFP. Genes deregulated in the four interconnected biological networks highlighted in Fig.4B are coloured in the same manner.

**Data file S1: Single-cell RNA-Seq data from the Malaria Cell Atlas.** Life stage-wise gene expression of 99 genes present in the four interconnected clusters shown in Fig. 4B based on *Plasmodium berghei* single cell RNA-seq data.

**Movies S1 to S2: Representative examples of ookinete motility in WT-GFP (S1) and Δ*mre11* (S2) lines.**
