## Supplementary figures and images for "MRE11 is crucial for malaria transmission and its absence affects expression of interconnected networks of key genes essential for life"

### Data file S1

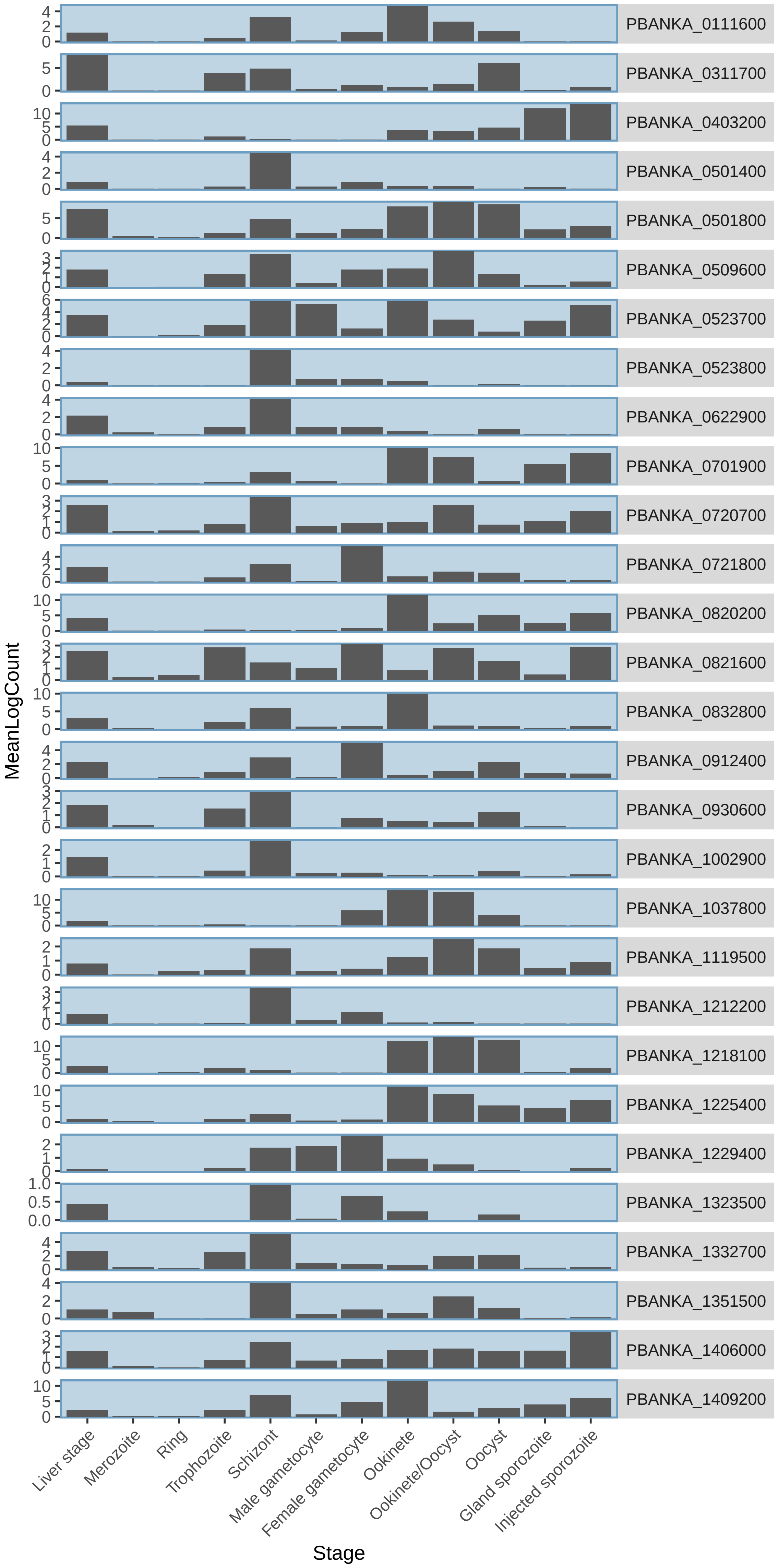

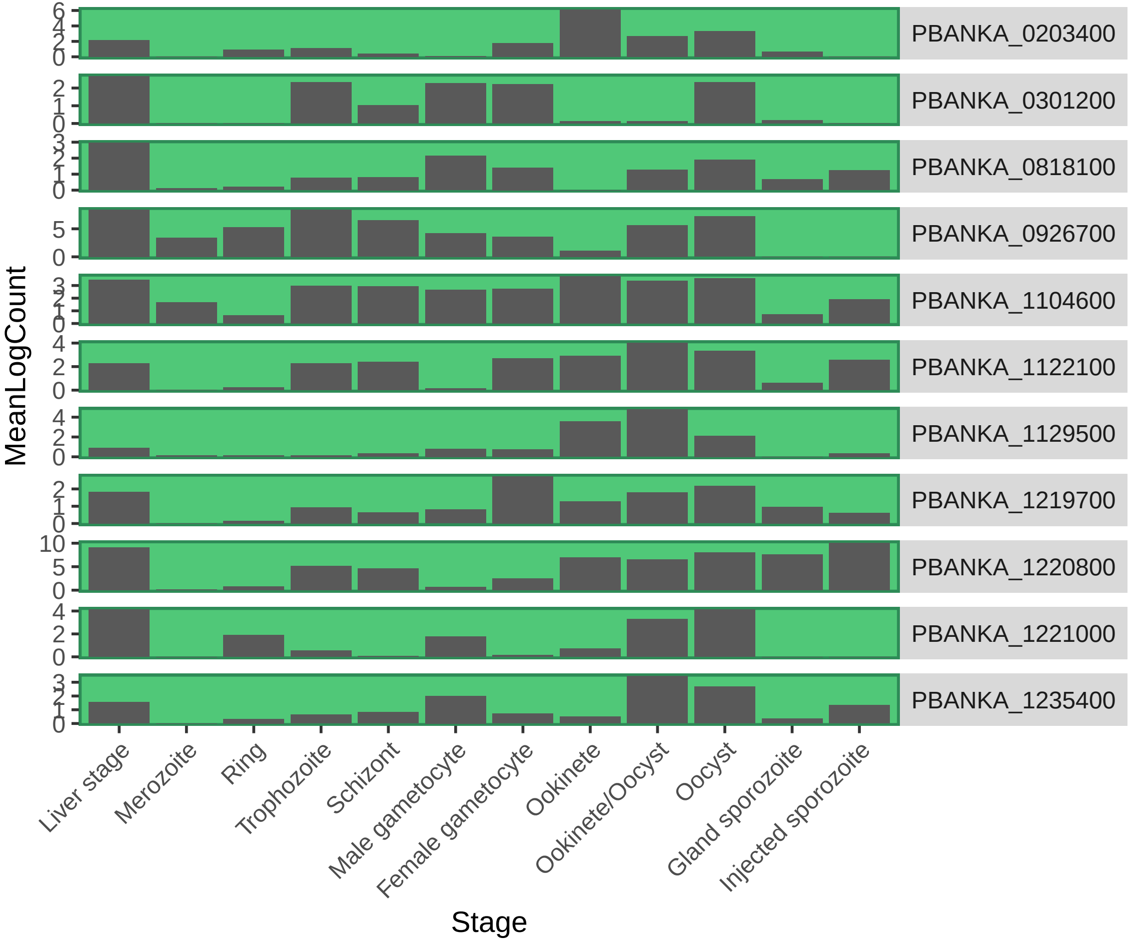

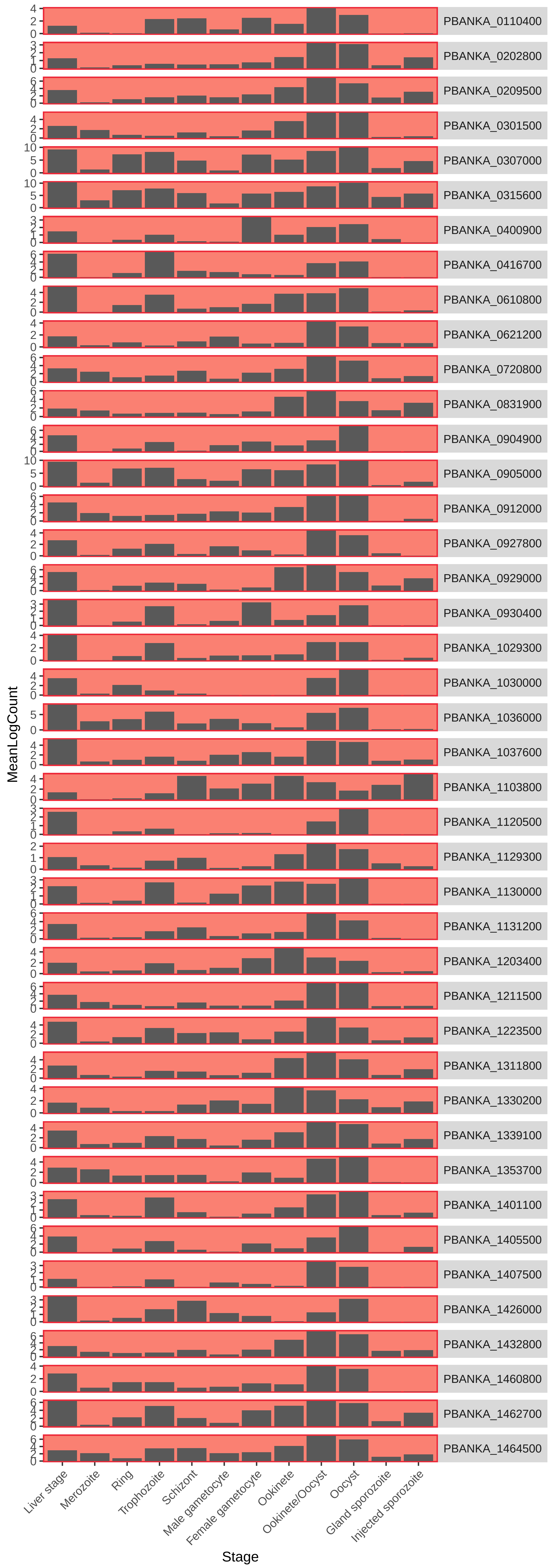

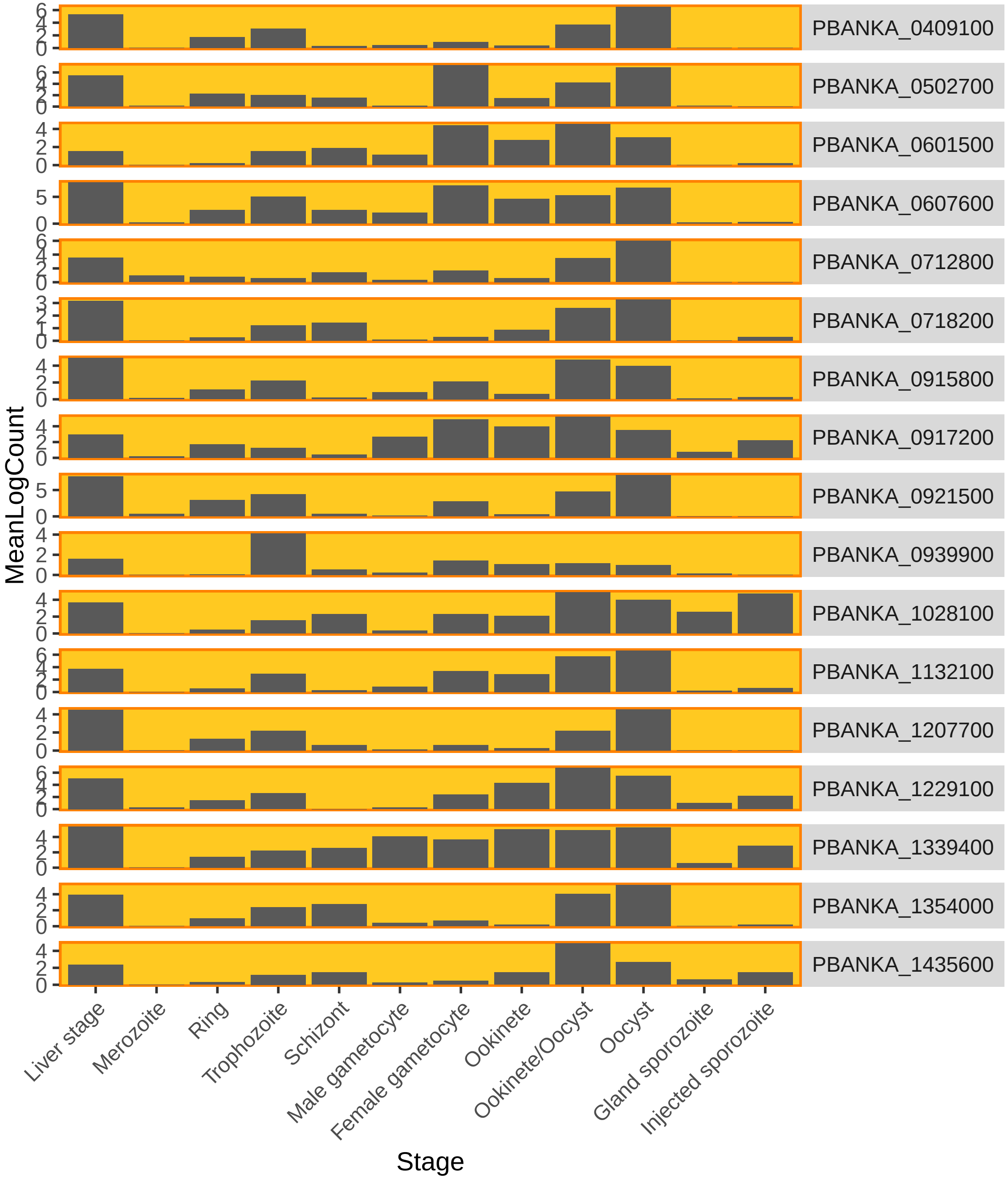
